# Ultrafast population coding and axo-somatic compartmentalization

**DOI:** 10.1101/2021.07.16.452655

**Authors:** Chenfei Zhang, David Hofmann, Andreas Neef, Fred Wolf

## Abstract

Populations of cortical neurons respond to common input within a millisecond. Morphological features and active ion channel properties were suggested to contribute to this astonishing processing speed. Here we report an exhaustive study of ultrafast population coding for varying axon initial segment (AIS) location, soma size, and axonal current properties. In particular, we studied their impact on two experimentally observed features 1) precise action potential timing, manifested in a wide-bandwidth dynamic gain, and 2) high-frequency boost under slowly fluctuating correlated input. While the density of axonal channels and their distance from the soma had a very small impact on bandwidth, it could be moderately improved by increasing soma size. When the voltage sensitivity of axonal currents was increased we observed ultrafast coding and high-frequency boost. We conclude that these computationally relevant features are strongly dependent on axonal ion channels’ voltage sensitivity, but not their number or exact location. We point out that ion channel properties, unlike dendrite size, can undergo rapid physiological modification, suggesting that the temporal accuracy of neuronal population encoding could be dynamically regulated. Our results are in line with recent experimental findings in AIS pathologies and establish a framework to study structure-function relations in AIS molecular design.

## Introduction

Humans, monkeys, and other mammals can perform complicated cognitive tasks that engage deep cortical hierarchies with processing times as brief a few hundred milliseconds ***Thorpe et al. (1996***); ***Stanford et al. (2010***). To achieve its overall speed, such fast information processing presumably depends on an ultrafast response dynamics of local cortical populations. Over the past years, numerous experimental studies tested different populations of cortical neurons for their capability to perform ultrafast population coding ***Carandini et al. (1996***); ***Köndgen et al. (2008***); ***Boucsein et al. (2009***); ***Higgs and Spain (2011***); ***Tchumatchenko et al. (2011***); ***Broicher et al. (2012***); ***Ilin et al. (2013***); ***Testa-Silva et al. (2014***); ***Ostojic et al. (2015***); ***Doose et al. (2016***); ***Nikitin et al. (2017***); ***Lazarov et al. (2018***); ***Linaro et al. (2018***). These studies used time domain and frequency domain analyses to assess the dynamics of the population’s mean firing rate in response to a shared input superimposed on a continuously fluctuating background, mimicking ongoing synaptic input. Using a time domain approach, it was shown for instance that populations of layer 2/3 pyramidal neurons in visual cortex nearly instantaneously respond to a small step in the average input ***Tchumatchenko et al. (2011***). In the frequency domain, the capability to generate an ultrafast response can be characterized by the population’s dynamic gain function. Dynamic gain was introduced theoretically by Bruce Knight in 1972 ***Knight*** (**1972a**) and measures the dynamic population rate response to sinusoidal input signals embedded in a background of i.i.d. statistically stationary noise. The frequency-domain correspondence of a sub-millisecond, ultrafast response is a dynamic gain function that starts to decay only for input frequencies above several hundred Hz. In cortical pyramidal cells, dynamic gain was first measured for populations of layer 5 pyramidal neurons, which indeed display a near-constant response up to a few hundred Hz ***Köndgen et al. (2008***), see also ***Carandini et al. (1996***) for an earlier indication of high bandwidth. As experimentally assessed, the dynamic gain function and the ultrafast response are not fixed and invariant properties of a neural population but substantially depend on the nature of background activity. Confirming a seminal theoretical prediction ***Brunel et al. (2001***), when the correlation time of the background input is increased, the dynamic gain in the high frequency regime is substantially boosted ***Tchumatchenko et al. (2011***), a phenomenon previously dubbed the Brunel effect ***Tchumatchenko et al. (2011***).

At first sight, the generation of ultrafast responses by an asynchronous population of neurons and its dependence on background fluctuations appear easy to understand. In the absence of external input, the cells fire asynchronously, because their background inputs are not correlated. This asynchrony guarantees that at any moment some fraction of the population is close to threshold and thus is in principle capable of an immediate, near instantaneous response. Intriguingly, however, several experimental studies demonstrated that a population’s propensity to generate an ultrafast response sensitively depends on subtle aspects of the cells’ biophysical and molecular design. In particular the molecular architecture of the spike initiation zone in the axon initial segment (AIS) seems to be specifically tailored to support an ultra-fast response. Lazarov et al. recently demonstrated that mutation of AIS-specific cytoskeletal proteins specifically impairs population response speed even if the ability of the cells to generate spikes per se is essentially unaffected ***Lazarov et al. (2018***). Consistent with this observation, the bandwidth of pyramidal neurons is substantially reduced after recovery from pathological conditions, such as transient hypoxia that affect AIS molecular architecture ***Revah et al. (2019***). Both of these findings are in line with prior studies suggesting that immature AIS organization ***Nikitin et al. (2017***) or pharmacological blockade of sodium channels at the AIS ***Ilin et al. (2013***) disrupt dynamic gain bandwidth and ultrafast population response. While the cellular and biophysical features sufficient to equip a cortical population with an ultrafast response are not completely understood, the weight of experimental evidence strongly suggests that this population-level property is sensitive to slight alterations in single neuron biophysics. It is interesting to note that phylogenetic studies place the response-speed-related refinements of AIS architecture at the invention of forebrain population coding in the first large chordate brains ***Lazarov et al. (2018***); ***Jenkins et al. (2015***); ***Hill et al. (2008***). A series of intriguing studies ***?Mohan et al. (2015***); ***Eyal et al. (2016***); ***Goriounova et al. (2018***) have even implicated inter-individual differences in the capability of human cortical neurons to support ultrafast population encoding as a determinant of heritable inter-individual difference in cognitive performance ***Goriounova et al. (2018***).

The feasibility of ultrafast population coding, its dependence on the structure of background input correlations, and its sensitivity to single neuron biophysical and molecular design have been correctly predicted by theoretical work preceding this recent wave of experimental studies. Theoretical studies, in single-compartment models, showed that ultrafast population encoding is closely associated with active AP initiation dynamics ***Fourcaud-Trocmé et al. (2003***); ***Naundorf et al. (2005)**; **Fourcaud-Trocmé and Brunel (2005***); ***Naundorf et al. (2006***); ***Wei and Wolf (2011***); ***Huang et al. (2012***); ***Ilin et al. (2013***). Neuron models that model AP generation by threshold-crossing, tacitly assuming an infinitely fast AP initiation, were first shown to exhibit a dynamic gain function with high bandwidth and the Brunel effect ***Brunel et al. (2001***). In subsequent studies ***Fourcaud-Trocmé et al. (2003***); ***Naundorf et al. (2005***); ***Fourcaud-Trocmé and Brunel (2005***), the voltage dependent sodium currents underlying AP initiation were incorporated. High bandwidth encoding could only be recovered when the voltage sensitivity of sodium currents was very high ***Fourcaud-Trocmé et al. (2003***). Based on these observations, the properties of the ion channels, which control the rapidness of AP initiation, were hypothesized as key parameters determining the bandwidth of the dynamic gain function ***Naundorf et al. (2006, 2007***).

The experimental studies summarized above suggest such a dependence in two ways. Firstly, the onset rapidness of the somatic AP waveform and the dynamic gain bandwidth measured in individual cells were found to be positively correlated ***Lazarov et al. (2018***); ***Nikitin et al. (2017***). Secondly, when the sodium channels at the AIS were blocked by TTX ***Ilin et al. (2013***) or when the number of ion channels at the AIS was reduced by genetic manipulation ***Lazarov et al. (2018***), the bandwidth of the dynamic gain function and the AP onset rapidness are both reduced. A distinct but related mechanism has been suggested to operate in multi-compartment models that describe the spatio-temporal dynamics of AP generation along the somato-axonal axis. APs are generated in the AIS, spatially separated from the soma, and lateral currents between these compartments are important for the detailed dynamics of AP initiation. Brette and coworkers ***Brette (2013***); ***Telenczuk et al. (2017)***, formulated an idealized multi-compartment model to study the impact of AP initiation site on somatic AP waveform. Increasing the distance between AP initiation site and soma, they predicted a discontinuous increase of the lateral current to emerge during AP generation. Even with a low intrinsic voltage sensitivity of the sodium currents underlying AP initiation, they suggested the conditions for an ultrafast population response might be fulfilled. A third hypothesis focuses on the impact of single neuron dendritic trees on population encoding ***Eyal et al. (2014***, ***2018***); ***Ostojic et al. (2015***). Eyal et al. ***Eyal et al. (2014***) first showed that multi-compartment models of the entire dendrite-soma-axon axis can realize high bandwidth encoding with a cutoff frequency above 100Hz, in particular in the presence of a large dendritic compartment. Intriguing support for this specific mechanism was found in a modeling and experimental study of Purkinje cells that exhibit an exceptionally large dendritic tree and show a very high bandwidth population response ***Ostojic et al. (2015***). All of these biophysical mechanisms, only a subset, or additional mechanisms yet unknown may underlie the presence or absence of ultrafast response dynamics in different neuronal populations. Irrespective of which of these alternatives ultimately apply, the increasing specificity with which recent experimental studies are probing the biophysical basis of ultrafast population coding calls for a systematic and controlled methodology to examine neuronal design variants at equivalent operating points and analyze the resulting impact on dynamic gain functions.

Here, we introduce such a methodology and use it to provide a dynamic gain analysis of the idealized multi-compartment model established by Brette ***Brette (2013***). In particular, we examine the impact on population response dynamics of various passive and active biophysical parameters representing AIS structure. To precisely compare the dynamic gain functions of different model variants at equivalent operating points, the mean and standard deviation of the inputs were chosen to maintain fixed average firing rate and coefficient of variation (CV) of the inter-spike interval (ISI) distributions. Stimulus filtering between the soma and the AP initiation site and the contribution of lateral currents to AP initiation are two fundamental differences between Brette’s model and simpler single compartment models. We find that electrotonic separation between AIS and soma and stimulus filtering does not, per se, result in the emergence of an ultrafast population response. Increasing the voltage sensitivity of sodium current induces high bandwidth and a sensitivity of dynamic gain to the input correlation (Brunel effect), demonstrating that this simple model is in principle capable of ultrafast encoding. In contrast, increasing the sodium conductance had no substantial impact on the dynamic gain function. Our results show that, in this idealized model, the somatic AP waveform has little relation to the emergence of an ultrafast population response. In contrast, the key determinants of ultrafast population coding are the voltage dependence of axonal sodium currents and the AP waveform at the AP initiation site around the time of AP onset.

## Results

### Voltage decoupling between soma and axon under voltage clamp and current clamp

We investigated the multi-compartment model introduced in ***Brette (2013***) (Fig 1 **A**) implemented in NEURON (see *Material and Methods*). Our implementation reproduced the central feature reported by Brette ***Brette (2013***): when the somatic voltage is clamped and changed so slowly that all currents attain their steady-state values, then we can observe a bifurcation in the in the voltage difference between soma and axon (dashed line in Fig 1**C**). The mechanism behind this bifurcation is explained Fig 1**B**. If the somatic voltage is fixed, the model is in steady state, if the sodium current equals the lateral current. This corresponds to the intersection points between the sodium current and the lateral current as a function of axonal voltage. The total resistance between the soma and the position of the sodium channels increases with *x*_Na_ and shapes the slope of the lateral current function. Increasing the clamped somatic voltage from −60mV to −50mV smoothly changes the intersection point of the two functions. However, when the position of sodium channels from the soma exceeds a critical distance, about 27*μ*m (Fig 1**B** middle panel), the number of intersection points increases to three. Thus the positioning initiation site away from the soma leads to a bifurcation of the equilibrium voltage as a function of somatic voltage. The equilibrium axonal voltage changes discontinuously and exhibits hysteresis. This results in an abrupt change of the lateral current entering the soma once the somatic voltage crosses a threshold value. This effect is a candidate mechanism to generate sharp APs at the soma and was hypothesized to induce ultrafast population encoding in cortical neurons ***Brette (2013***).

**Figure 1.**
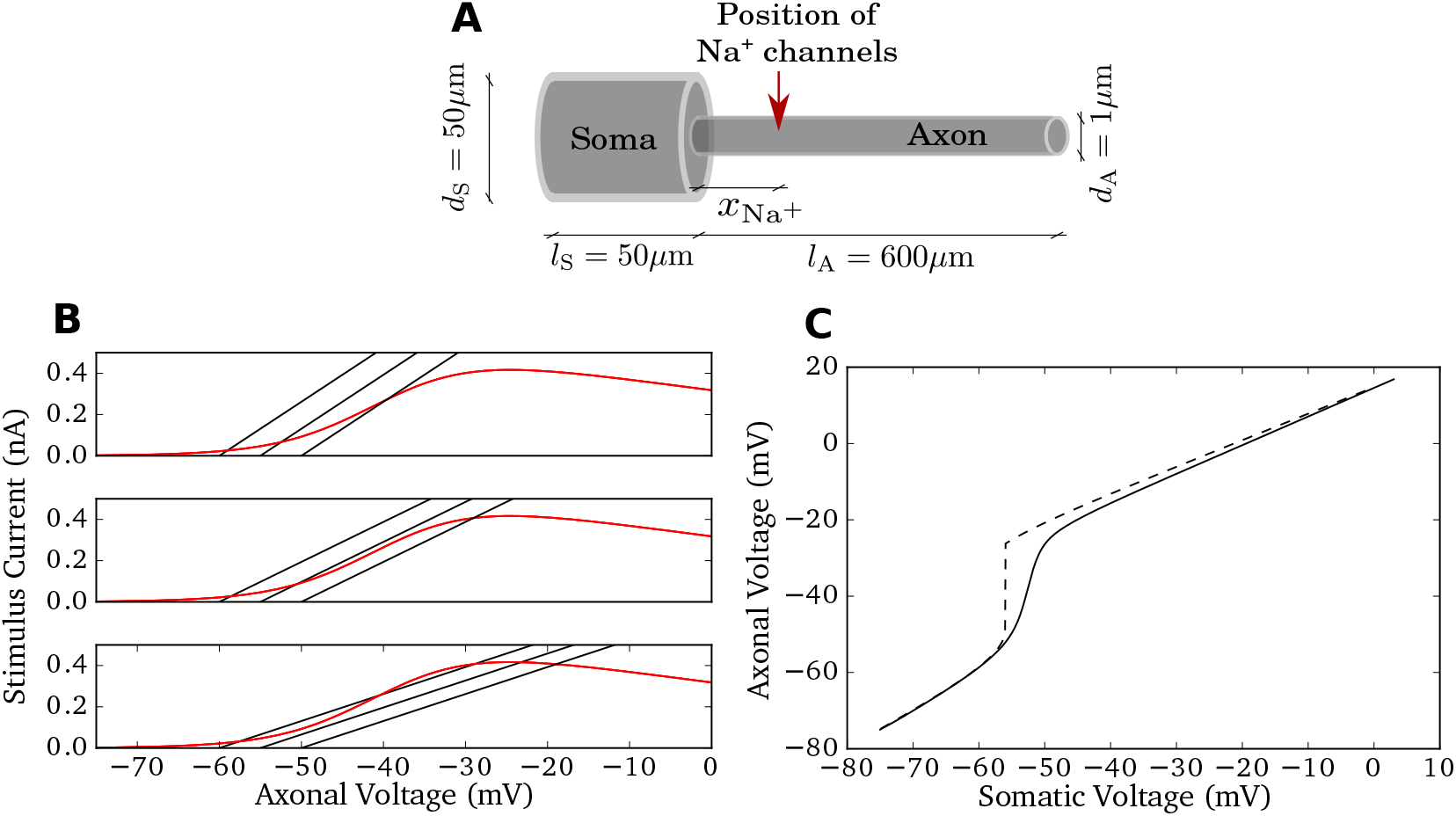
Model properties. **A** Sketch of the model morphology: soma and axon are modeled as cylinders. The sodium conductance is positioned at *x*_Na_ and the stimulus current is injected at the middle of the soma. For simulation and model details see methods. **B** Bifurcation of axonal voltage in compare Fig 2 of ***Brette (2013***). Sodium current (red) and lateral currents (black, with somatic voltage fixed at −60mV, −55mV, −50mV respectively) are plotted as a function of axonal voltage at different *x*_Na_ (top to bottom: *x*_Na_ = *20μm, 27 μm*, 40*μ*m). The intersection points between the black curve and the red curve indicate stationary axonal voltage given the somatic voltage. For large *x*_Na_ = 40*μ*m, the first intersection point changes discontinuously for increasing somatic voltage. **C** Axonal voltage as a function of somatic voltage under voltage clamp (dashed) and current clamp (solid) at the soma for *x*_Na_ = 40*μ*m. For current clamp, the soma is injected with a constant current that generates 5Hz firing rate. The voltage clamped case shows a loss of voltage control around −55mV. The dynamic case, however, shows a less sharp transition.

In Fig 1**C**, we plot the axonal voltage as a function of somatic voltage for *x*_Na_ = 40*μ*m. With the somatic voltage clamped to different values, we recorded the corresponding stationary axonal voltages. As expected, there is a discontinuity around the somatic voltage of −55mV, the somatic voltage at which two intersections between lateral and sodium current meet. This corresponds to the annihilation of a stable and an instable fixpoint. Apart from simulations with a voltage clamped soma, we also studied the current clamp condition, in which we inject a slowly increasing input current in the middle of the soma, and recorded the two voltage traces at the soma and at the AP initiation site. In this case, the lateral current from axon to soma not only slows axonal depolarization, but accelerates the depolarization of the soma. As shown in Fig 1**C** (continuous line), the decoupling of somatic and axonal voltages was substantially smoother in this dynamic case. Even if the parameter *x*_Na_ is sufficiently large for the bifurcation under the voltage clamp case, we only observed a gradual deviation of the two voltages. The deviation speed is limited by the activation speed of the sodium current. The maximum amplitude of the sodium current remained unaffected, so for larger voltages, the decoupling distances of the two voltages are maintained.

### Spatial decoupling is not sufficient for ultrafast population encoding

Next, we characterized the sharpness of the somatic action potential as we varied the sodium channel position *x*_Na_ from 0*μ*m to 80*μ*m (Fig 2**A**). We found that the somatic voltage rises more rapidly, the further away from the soma the sodium channels were positioned. The AP initiation dynamics at *x*_Na_ = 27*μ*m did not exhibit a bifurcation.

**Figure 2.**
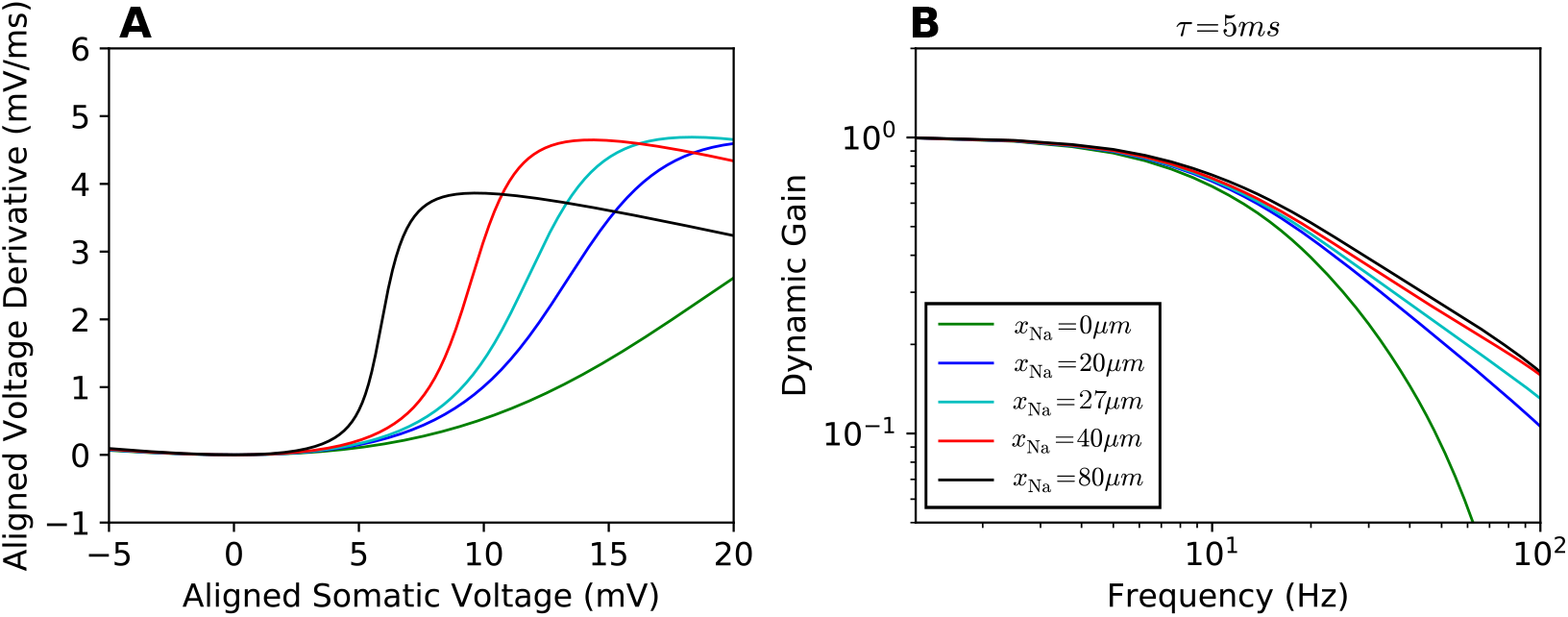
Phase plot and linear response functions. **A** Phase plots for the APs seen at the soma, somatic voltage rate of change vs. somatic voltage for different sodium channel positions *x*_Na_ (legend in **B**). The local minima of all curves are aligned at (0mV, 0mV/ms). **B** Dynamic gain functions for different *x*_Na_, normalized. 95% confidence intervals are plotted as shaded areas, mostly hidden by the line width. All curves are above statistical significance threshold (see *Materials and Methods*). The input current correlation time was set to *τ* = 5ms.

We then calculated the dynamic gain for various values of the position parameter *x*_Na_ as depicted in Fig 2**B**. For these calculations, we fixed the firing rate and CV as described in methods, and calculated different dynamic gain curves for the *x*_Na_ values. The correlation time of the fluctuating background current was set to 5ms. Moving the sodium channels away from the soma somewhat enhanced the high frequency response, in particular for *x*_Na_ = 0 to *x*_Na_ = 20*μ*m. The distance for which a bifurcation in the voltage-clamp condition was observed does not stand out in any way. Moreover, all obtained dynamic gain curves exhibited a cut-off frequency of about 10Hz and the dynamic gain curves converged in a form similar to a canonical conductance-based neuron model with a decay exponent of −1 ***Fourcaud-Trocmé et al. (2003***). Increasing *x*_Na_ from 40*μ*m to 80*μ*m had limited effect on the dynamic gain curve, despite doubling the electrotonic separation between AP initiation site and soma.

Compared to a single compartment model, there are two main differences in fluctuation-driven AP initiation for the current multi-compartment model. Firstly, in a single compartment model, the injected stimulus waveform directly enters the dynamic equation of AP generation. In the multi-compartment model, the spatial separation between injection site and initiation site creates a filter. The cable properties attenuate the stimulus as it is transmitted to the AP initiation site. This may decrease the dynamic gain in the high frequency range relative to a single compartment model. Secondly, the current components underlying AP generation are modified. In a single compartment model the entire local sodium current is effective for AP initiation. In the multi-compartment model, the sodium current, entering at the AP initiation site, not only charges the local membrane capacitance, but also feeds lateral currents that first depolarize neighboring parts of the axon and subsequently the somatic membrane. This lateral current from the AP initiation site depends not only on the distance to the soma, but also on the axial resistivity of the axon. Thus the axial resistivity is an additional parameter that impacts on the voltage dynamics at the AP initiation site of a multi-compartment model.

The lateral current away from the AP initiation site slows down the dynamics of AP generation. Moving the AP initiation site away from the soma (Fig 2**A**), we found that the shape of the APs at the soma is changed, but the AP initiation dynamics in the axon is modified much more mildly. Fig 3**A** and **B** show the impact of lateral current and electrotonic filtering on axonal voltage dynamics and population encoding. In Fig 3**A**, the phase plot for APs at the AP initiation site shows that shifting the AP initiation site away from the soma increases the resistance between, thus reduces the lateral current towards the soma during the AP upstroke. For larger *x*_Na_ values, the AP initiation dynamics thus becomes more rapid. Fig 3**B** depicts the filtering effect due to stimulus transmission from the soma to the AP initiation site. We injected the soma with the same stimuli that were used for calculating dynamic gain functions. We used the axonal voltage at *x*_Na_ as the continuous response variable and calculated the transfer function from somatic input current to axonal voltage at *x*_Na_. The dominant effect is the filtering from current to somatic voltage. With increasing *x*_Na_ a small additional damping of high frequency components appeared. Below 100Hz, this additional damping effect was weak for *x*_Na_ = 20*μ*m, 40*μ*m, 80*μ*m. These results suggest that the dynamic gain values are enhanced for larger *x*_Na_ because the larger axonal resistance reduces the lateral current while the input experiences only minor damping. Further increasing *x*_Na_, one expects that the AP initiation dynamics saturates while the electrotonic filtering will eventually reduce the high frequency gain.

**Figure 3.**
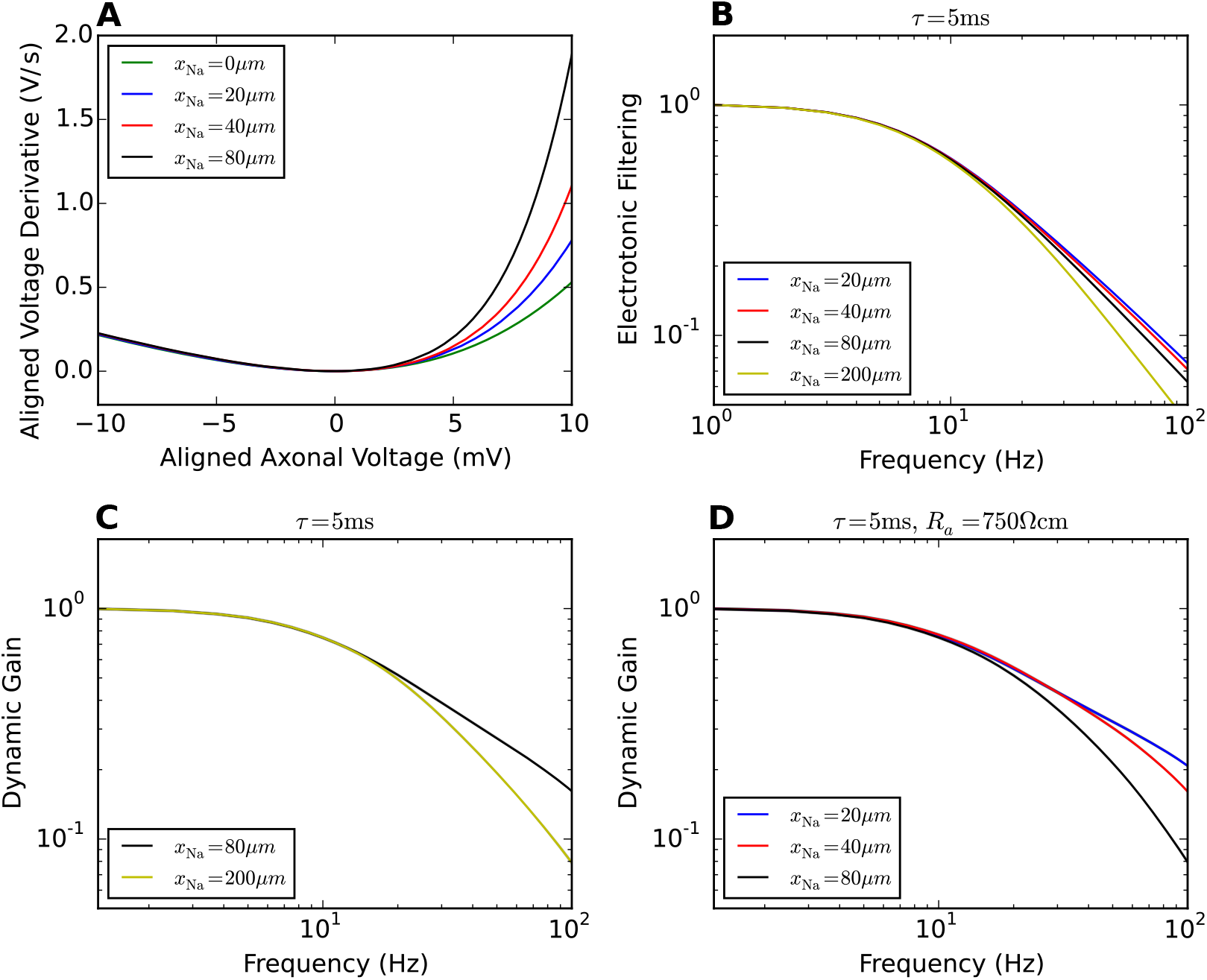
Impact of lateral current and electrotonic filtering on dynamic gain. **A** Phase plots for the AP waveforms at the initiation site. Moving the initiation site away from the soma reduces the lateral current during AP generation, leading to a more rapid initiation dynamics. The local minima of all curves are aligned at (0mV, 0mV/ms). **B** Electrotonic Altering when the stimulus is transmitted from the soma to the AP initiation site. **C** Increasing *x*_Na_ to 200*μ*m decreases the high frequency gain of the neuron model. **D** Increasing *R_a_* to 750Ωcm reduces the impact of lateral current on dynamic gain. The electrotonic filtering effect dominates the dynamic gain. A larger *x*_Na_ reduces the high frequency gain.

This should lead to an impaired encoding ability for high frequencies. Fig 3**C** confirms this prediction. We calculated the dynamic gain function for *x*_Na_ = 200*μ*m. The dynamic gain was reduced compared to that for *x*_Na_ = 80*μ*m. Another expectation is that by reducing the lateral current, the dynamic gain functions should become more sensitive to electrotonic filtering with larger *x*_Na_. To examine this, we increased the axial resistance *R_a_* fivefold to 750Ωcm. The dynamic gain functions in Fig 3**D** show that with lateral current suppressed, increasing *x*_Na_ reduces the high frequency dynamic gain monotonically opposite to the behaviour shown in Fig 2**B**.

### Negligible impact of sodium peak conductance on dynamic gain

Experimental studies have shown that a high density of sodium channels in the AIS is required for high bandwidth encoding ***Ilin et al. (2013***); ***Lazarov et al. (2018***). We thus investigated how changing sodium peak conductance impacts on population encoding in the multi-compartment model. The sodium peak conductance was increased 5 and 10 fold. Fig 4**A** and **C** shows the dynamic gain curves for 1Hz and 5 Hz firing rate. The CV was controlled to vary less than 0.1 for both rates. Increasing the sodium peak conductance caused only minor changes to the dynamic gain curves. For the 1 Hz firing rate, the dynamic gain at high frequencies was slightly enhanced for higher sodium peak conductance. Further increasing the sodium peak conductance to 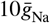, the gain curve remained almost identical to that at 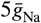. For 5Hz firing rate the encoding of high frequencies was overall better than for the 1 Hz condition. However, increasing the sodium peak conductance lead to a slight decrease of dynamic gain at high frequencies. The main effect of increasing sodium peak conductance in the model was a substantial lowering of the current and voltage threshold of the model. However, static gain around threshold, i.e. the slope of the F-I curves, was largely preserved. The inset in panel **C** shows the F-I curves for the three sodium peak conductances used. Note that the F-I curves are shifted along the current axis. This essentially preserves gain as measured by the curve’s slope at a fixed firing rate.

**Figure 4.**
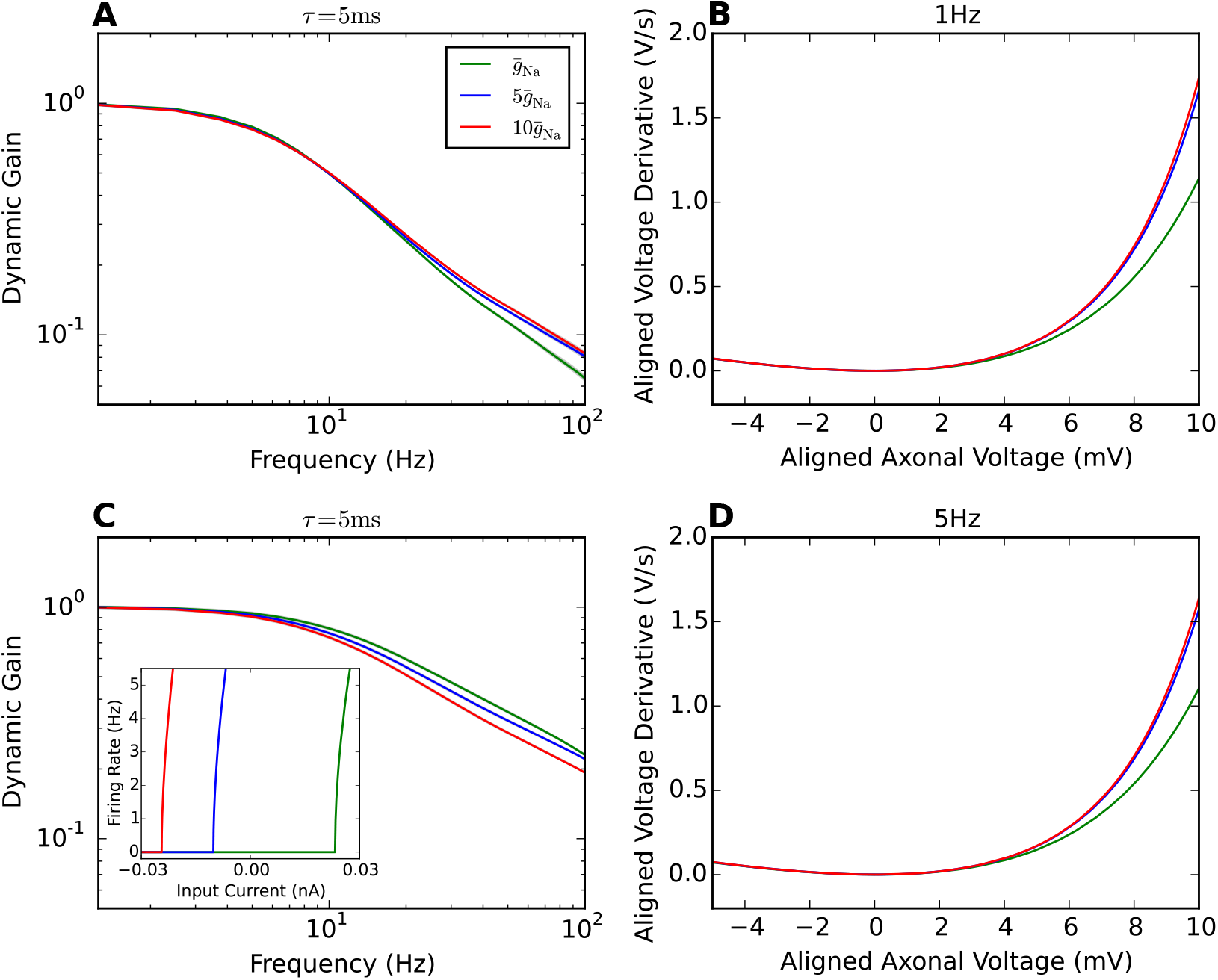
AP initiation dynamics and dynamic gain comparisons for difference sodium peak conductances. **A** Dynamic gain functions of neuron models with different sodium peak conductances for 1Hz firing. The CV is 0.9±0.05. **B** Phase plots of AP initiation dynamics of Brette’s models with different sodium peak conductances. *x_Na_* is fixed to 40*μ*m. Sodium peak conductance 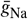 is increased 5 and 10 fold separately in comparison with the original model. To compare the AP initiation speed, each neuron model is injected with a constant input that generates 1Hz firing rate. **C** Dynamic gain functions of neuron models with different sodium peak conductances for 5Hz firing. The CV is 0.85±0.05. The inset in panel **C** shows the F-I curve for the three sodium peak conductance used. **D** Phase plots of AP initiation dynamics at 5Hz firing.

Fig 4**B** and **D** show the phase plots of AP initiation dynamics for constant stimuli that reproduce 1Hz and 5Hz firing rate separately. In both conditions, increasing the sodium peak conductance by a factor of 5 accelerated AP initiation. A further doubling had less impact in the voltage range displayed here. Comparing the AP initiation dynamics with the dynamic gain functions, at 1Hz firing rate, the dynamic gain enhancement parallels the AP upstroke acceleration for increasing sodium peak conductance. For the 5Hz firing rate, the small changes in dynamic gain at 100 Hz appear dissociated from the changes of AP dynamics.

We next combined the variation of initiation site location and sodium peak conductance. The resulting cutoff frequencies of the dynamic gain are displayed in Fig 5 for at fixed firing rates of 5Hz and 1Hz. The exact working points were found by fixing the the CVs of the ISI distributions to 0.9 ± 0.05 for Fig 5**A** and **B**, or by fixing the std of stimuli for Fig 5**C** and **D**. The 2D surface plots show the parameter dependency of the cutoff frequencies and the decay of dynamic gain values in the low frequency region. At 5Hz firing rate, both stimulus condition in **A** and **C** show that the cutoff frequency is higher with larger *x*_Na_ and smaller sodium peak conductance. At high sodium peak conductance, the cutoff frequency is less sensitive to the AP initiation site. At 1Hz firing rate, we observed a similar trend of the cutoff frequency. Compared to the high firing rate cases, at 1Hz firing rate, the 2D surface plots are flatter for both stimulus condition. Besides, the CV of the ISI distribution was less dependent on the mean at 1Hz firing rate. Although the dynamic gain values in the high frequency region are enhanced for higher sodium peak conductance at 1Hz firing rate, the cutoff frequencies are slightly lower for high sodium peak conductance models. When fixing the std, we obtained a similar dependence.

**Figure 5.**
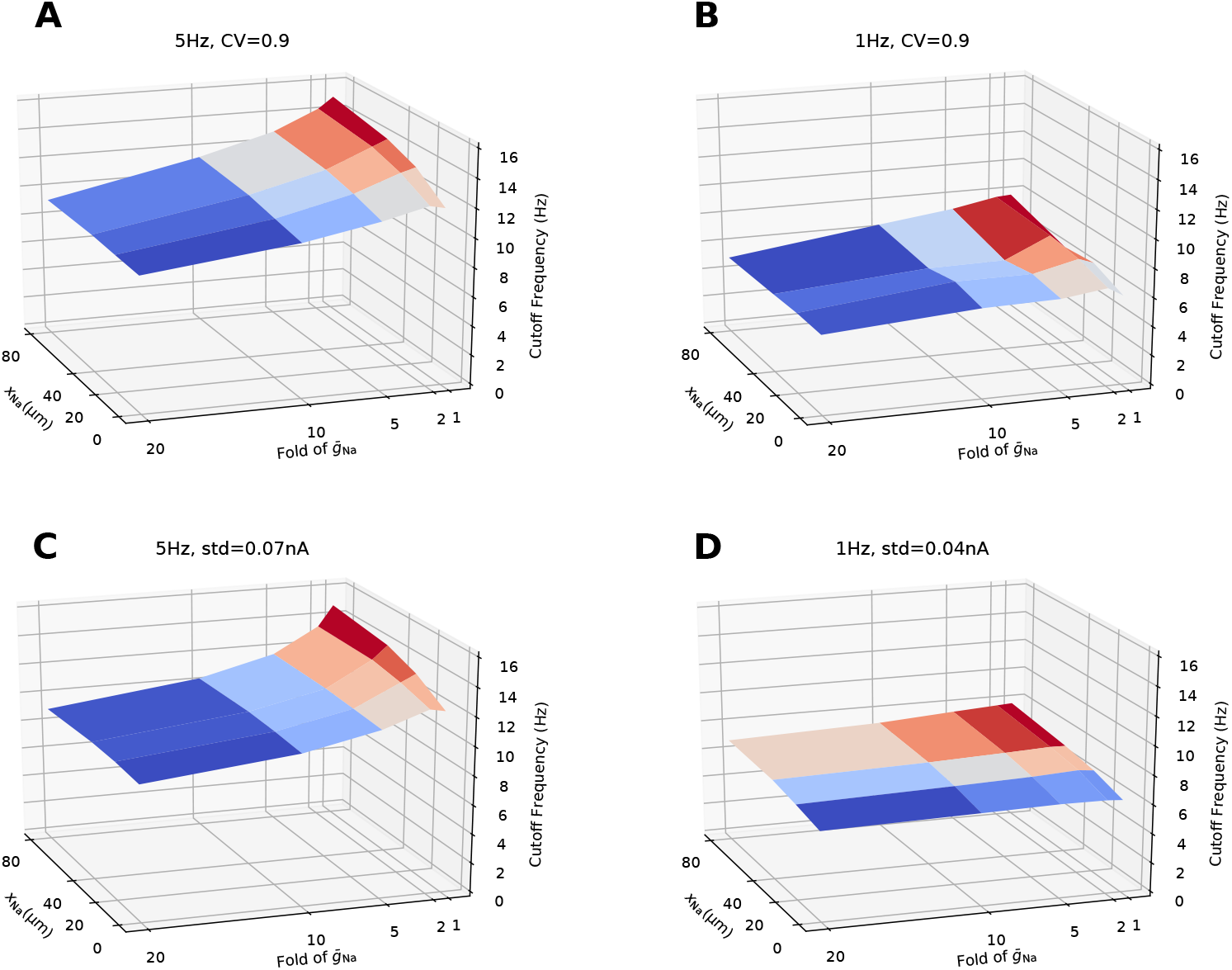
Cutoff frequencies of dynamic gain curves as a function of sodium peak conductance and AP initiation site. **A** and **B** For each pair of AP initiation site and sodium peak conductance, the dynamic gain curve was calculated at 5Hz and 1Hz firing rates working points. CV values were fixed at 0.9 ± 0.05. **C** and **D** The dynamic gain curves were calculated at 5Hz and 1Hz firing rate with std of stimuli fixed at 0.07nA and 0.04nA. Cutoff frequency was the frequency to which the dynamic gain decayed by a factor of 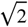. Correlation time of background current was 5ms. *x*_Na_ ranged from 0*μ*m to 80*μ*m. Sodium peak conductance ranged from 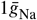 to 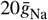.

### Weak high-frequency boost when axonal currents show moderate voltage sensitivity

Next, we examined the impact of correlation time of background current fluctuations on the dynamic gain curve (Fig 6). We increased it from *τ* = 5ms to 50ms and calculated the dynamic gain functions for *x*_Na_ = 20*μ*m, 40*μ*m, 80*μ*m. Distinct from the behaviour of LIF-like models ***Brunel et al. (2001***); ***Naundorf et al. (2005***); ***Tchumatchenko and Wolf (2011***); ***Puelma Touzel and Wolf (2015***); ***Fourcaud-Trocmé et al. (2003***) and cortical neurons ***Tchumatchenko et al. (2011***), increasing the correlation time did not enhance the dynamic gain in the high frequency range. We observed a weak resonance around 10Hz, i.e. a slight enhancement of dynamic gain below the cutoff frequency. For higher frequencies, the dynamic gain decayed steeper than that for *τ* = 5ms. This observation holds for all three *x*_Na_ values, and shows that overall the dynamic gain of the model is quite insensitive to input current correlation time. Independent of the input correlation time, this model’s dynamic gain had a much lower bandwidth than any of the experimentally characterized cortical neurons, which exhibit cutoff frequencies in the range of hundreds of *Hz**Ilin et al. (2013***); ***Köndgen et al. (2008***); ***Tchumatchenko et al. (2011***); ***Boucsein et al. (2009***); ***Higgs and Spain (2009***); **?**); ***Ostojic et al. (2015***); ***Linaro et al. (2018***); ***Broicher et al. (2012***); ***Lazarov et al. (2018***); ***Carandini et al. (1996***); ***Higgs and Spain (2011)**; **Doose et al. (2016***); ***Nikitin et al. (2017***); ***Volgushev (2016***). The insensitivity of the dynamic gain to the current correlation time qualitatively deviates from experimental observations ***Tchumatchenko et al. (2011***). In summary, our results show that shifting the position of the AP initiation site away from the soma does increase the onset rapidness of APs observed at the soma, but sharp somatic APs onset does not by itselfe induce an ultrafast population response.

**Figure 6.**
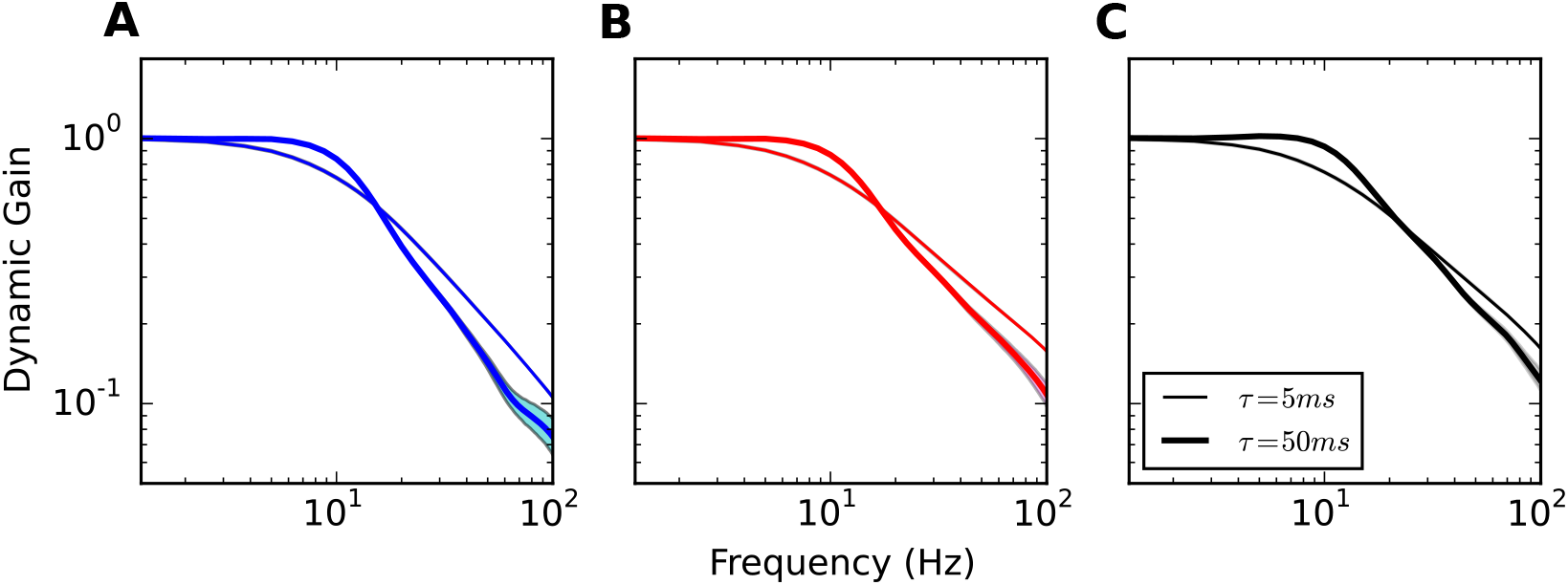
Impact of background current correlation time on dynamic gain. Dynamic gain functions for different input current correlation times. Model properties as for Fig 2. *x*_Na_ increases from = 20,40 to 80*μ*m (**A**, **B**, **C**). Line width indicates input correlation time (thin: *τ* = 5ms, wide: *τ* = 50ms).

### Increasing axonal current’s voltage sensitivity establishes ultrafast population encoding

The results above suggest that the AP voltage dynamics at the AP initiation site is more closely related to the high frequency encoding properties than the somatic AP waveform. We thus examined whether increasing the voltage sensitivity of the sodium current at the initiation site could enhance high frequency gain and potentially equip the multi-compartment model with an ultrafast population response.

To test this, we changed the parameter *k_a_* of the activation function of the sodium gating variable from 6mV to 0.1mV making the sodium current activation curve nearly step-like. With this modification, the voltage decoupling between soma and axon was nearly instantaneous for all *x*_Na_ tested (Fig 7**A**). With high voltage sensitivity, the axonal voltage and somatic voltage are nearly identical until *V*_1_/_2_ is reached. Above this threshold value, a large sodium current is activated, causing an immediate substantial decoupling. Compared to the voltage decoupling with a standard sodium activation function, there was a substantial change of the decoupling trajectory for *x*_Na_ = 20*μ*m. However, for large *x*_Na_ values, the decoupling trajectories are quite parallel with those for a less voltage sensitive sodium activation function. In Fig 7**B**, we compare the dynamic gain functions of original model to the model with more voltage sensitive sodium activation function. We fixed *x*_Na_ to 40*μ*m and *τ* to 5ms. Fig 7**B** shows that the dynamic gain decreases less for high frequencies and hence the bandwidth increases.

**Figure 7.**
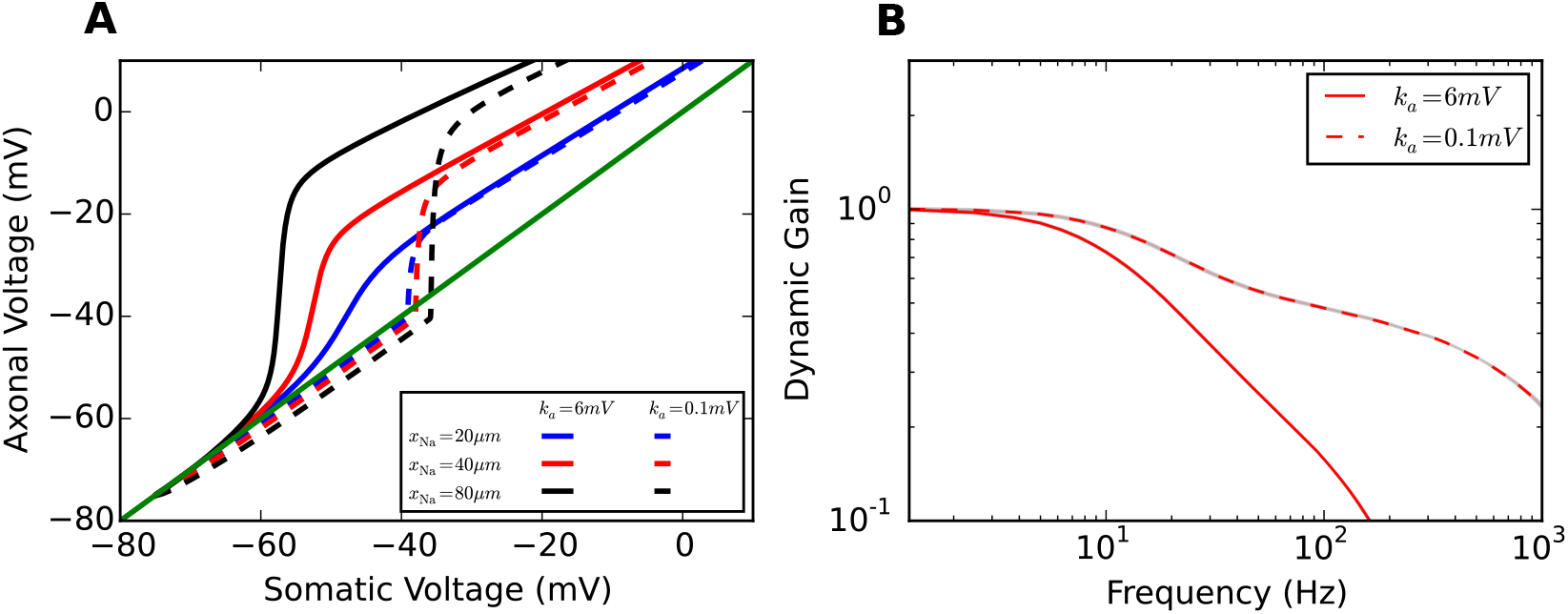
Impact of voltage sensitivity of sodium current activation curve on voltage decoupling under current clamp and dynamic gain. **A** Plot of axonal voltage at *x*_Na_ vs. somatic voltage for different sodium channel positions. Solid lines are voltage traces for low voltage sensitivity *k_a_* = 6mV, dashed lines denote the traces for high voltage sensitivity *k_a_* = 0.1mV. **B** Dynamic gain functions for the different voltage sensitivities for *x*N_a_ = 40*μ*m and *τ* = 5ms

We also examined the impact of the positioning of the AP initiation site and the input current correlation time on dynamic gain for the case of high voltage sensitivity (Fig 8). Fig 8**A** shows that with *k_a_* = 0.1mV, the high frequency dynamic gain is greatly enhanced for all three *x*_Na_ values. One additional observation is that increasing *x*_Na_ decreases the dynamic gain in the high-frequency regime. This behavior is opposite to the behavior for low voltage sensitivity (Fig 2**B**). This phenomenon may be related to the impact of electrotonic filtering and lateral current proposed above. When the sodium activation function is highly voltage sensitive, the lateral current has limited impact on the voltage dynamics at the initiation site. However, the electrotonic filtering from soma to axon still impacts the dynamic gain function in the high frequency range. Fig 8**B** displays the dynamic gain functions for *τ* = 50ms. For higher voltage sensitivity, increasing the input current correlation time increases high frequency gain similar to the behavior first described for the LIF neuron ***Brunel et al. (2001***).

**Figure 8.**
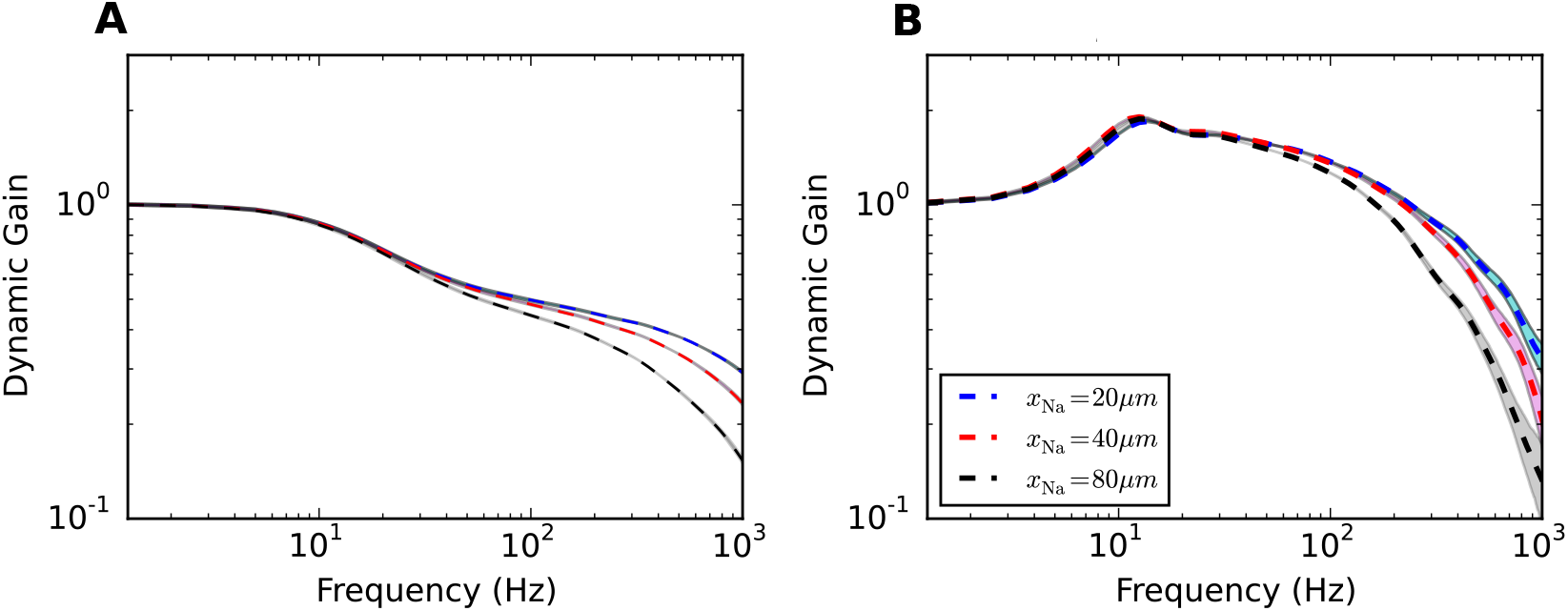
Increased impact of current correlation time for high voltage sensitivity. Dynamic gain curves are plotted for **A** a short input current correlation time *τ* = 5ms and **B** a long correlation time of *τ* = 50ms. Dashed lines as in Fig 7**B**. Shaded areas are 95% confidence intervals. Colors denote different initiation site positions (see legend).

The activation dynamics of sodium current during AP initiation is determined by both the voltage sensitivity of sodium activation curve *k_a_* and the sodium peak conductance 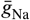. In Fig 9, we display the cutoff frequency as a function of these two parameters. We fixed *x*_Na_ at 40*μ*m and correlation time of background current *τ* at 50ms. *k_a_* ranged from 0.1 to 6mV. 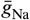 was increased from 1 fold to 20 fold. We fixed the firing rate and CV (Fig 9**A** and **B**), or firing rate and std of stimulus (Fig 9 **C** and **D**). For all four conditions, increasing the voltage sensitivity of sodium activation curve increased the cutoff frequency in agreement with previous theoretical studies ***Brunel et al. (2001***); ***Fourcaud-Trocmé et al. (2003***); ***Wei and Wolf (2011***); ***Huang et al. (2012***). The cutoff frequencies for 5Hz firing rate were in general larger than those for 1Hz firing rate at given *k_a_* and 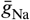. Overall, the cutoff frequencies were much less sensitive to 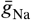 than to *k_a_*. Fig 5 shows that at *k_a_* = 6mV and *τ* = 5ms, the cutoff frequency is slightly decreased when increasing the sodium peak conductance. For *k_a_* = 6mV and *τ* = 50ms, a higher sodium peak conductance also led to a slightly lower cutoff frequency. For other values of *k_a_*, the cutoff frequency remained relatively insensitive to the sodium peak conductance.

**Figure 9.**
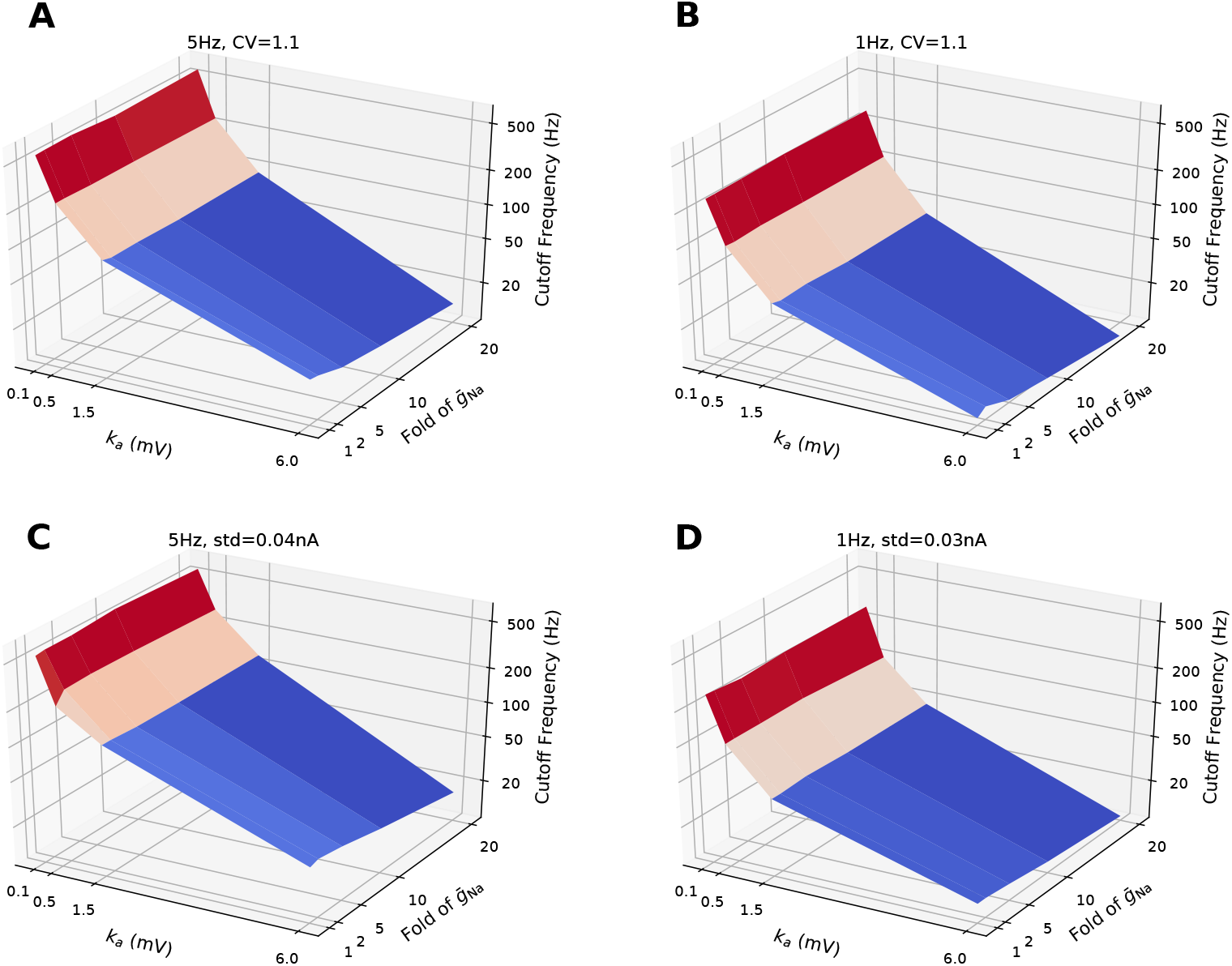
Cutoff frequencies of dynamic gain curves as a function of sodium peak conductance and voltage sensitivity of sodium activation curve. **A** and **B** For each pair of voltage sensitivity and sodium peak conductance, the dynamic gain curve was calculated at 5Hz and 1Hz firing rates working points. CV values were fixed at 1.1 ±0.05. **C** and **D** The dynamic gain curves were calculated at 5Hz and 1Hz firing rate with std of stimuli fixed at 0.04nA and 0.03nA. Correlation time of background current was 50ms. *x*_Na_ fixed at 40*μ*m. Sodium peak conductance ranged from 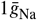 to 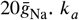 ranged from 0.1 to 6mV.

### Excessive soma size improves high-frequency dynamic gain

Quantitatively it is not clear what type of impact changing the soma size may have on dynamic gain. On the one hand, following the rationale originally laid out by Brette, increasing the size of the soma could be expected to make the dynamics of the model more representative of the voltage clamp behavior depicted in Fig 1. The larger the soma, the lower is the impact of the lateral currents on the somatic voltage and the more tightly is the voltage at the origin of the axon clamped to the somatic voltage. From this perspective, the model should effectively behave like a LIF model in the limit of a very large soma. On the other hand, if the membrane potential dynamics at the AP initiation site is predictive of dynamic gain, increasing the soma size may have the opposite effect. Then the larger the soma, the more current should be diverted to charge the soma slowing down the axonal voltage upstroke and reducing the high frequency dynamic gain. Brette’s original model ***Brette (2013***) had a soma size substantially larger than that of cortical neurons. To assess the effects sketched above and also determine the behavior at realistic soma size, we decided to vary *d_S_* and *l_S_* to examine the impact of soma size on dynamic gain.

In the following calculations, *d_S_* and *l_S_* ranged from 10*μ*m to 2cm. *x*_Na_ was 40*μ*m, and the input current correlation time was 5ms. For the 2cm case, the spatial grid of the soma was increased to 1000*μ*m for the convenience of simulation. As depicted in Fig 10**A**, the colored thick lines are the unnormalized dynamic gain curves for various soma sizes. To generate the same firing rate in these models, similar amounts of current reached the AP initiation site. However, to overcome the leak current in the soma and generate somatic voltage fluctuations, the somatic input had to be scaled with the surface area of the soma. As a result, the dynamic gain amplitudes were greatly attenuated for the larger soma cases.

**Figure 10.**
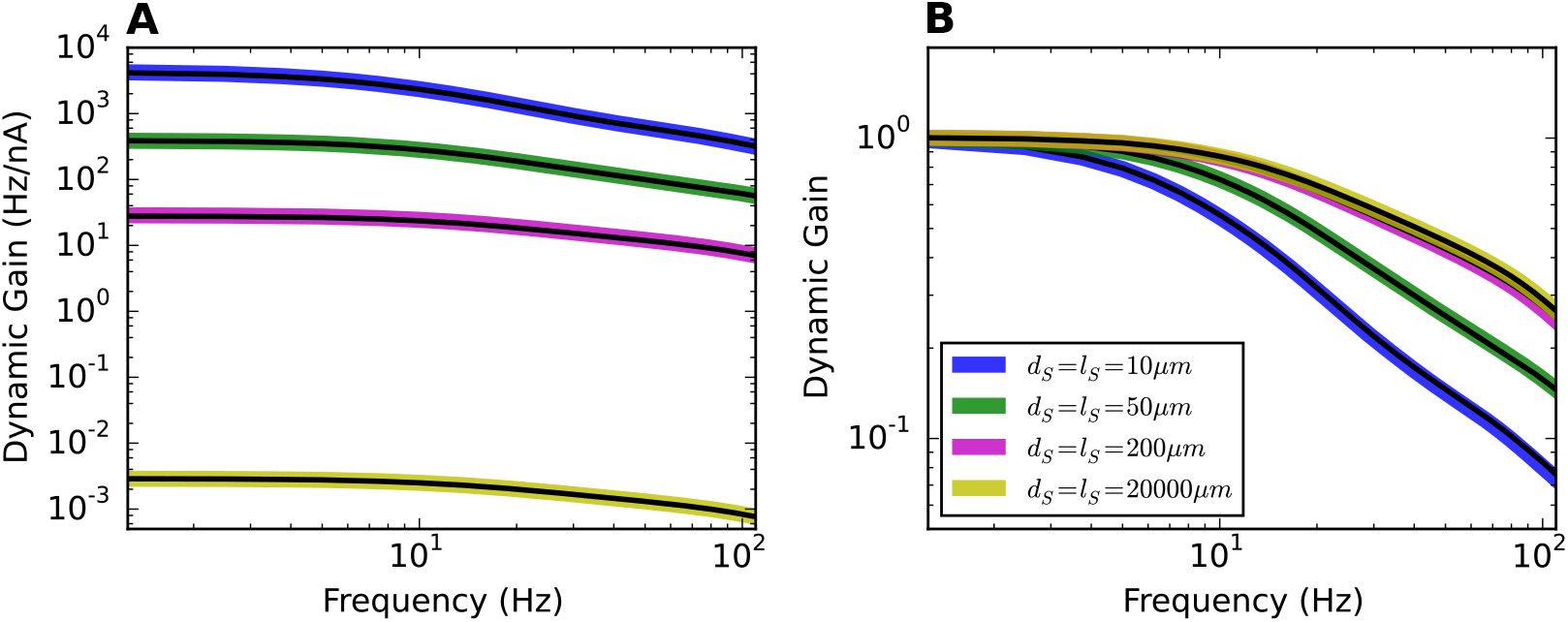
Effect of soma size on dynamic gain. **A** Unnormalized dynamic gain curves for various soma sizes. Diameters and lengths for all the soma are shown in the legend panel in **B**. *x_Na_* was fixed to 40*μm*. The correlation time *τ* was fixed to 5ms. For each soma size, fixing surface area but reducing soma length to 2*μ*m, the dynamic gain curves were recalculated and represented as black lines, which overlapped with corresponding colored lines. **B** Comparison of all the normalized dynamic gain curves. Increasing the soma size enhances the dynamic gain in the high frequency regime for Brette’s original model.

Although of “enormous” length in the most extreme case, the somatic compartment was still electrotonically compact in these simulations. To test that the electrotonic filtering was in fact minimal, we kept the soma surface areas and reduced the soma length to 2*μ*m and recalculated the corresponding dynamic gain curves. The black lines in Fig **10A** demonstrates that electrotonic filtering in the soma has negligible influence on the population dynamic gain.

In Fig 10**B**, we compared the normalized dynamic gain curves for different soma sizes. Increasing the soma size enhanced the high frequency dynamic gain. However, the dynamic gain curves did not become flat as expected for a LIF-like model. As a function of soma size, the normalized dynamic gain seemed to converge to a limiting curve, with little change beyond soma sizes of 200*μ*m. The deviation of the neuron model from the LIF model may be explained by the position of the AP initiation site. When the sodium channels are positioned at the soma, for a given sodium peak conductance, increasing the soma size decreases the ability of sodium current to change the voltage derivative. In the large soma size limit, the sodium current would not suffice to overcome the leak current for AP generation, so that the neuron model will be closely approximated by a model with hard threshold driven only by the external input, i.e. a LIF model. However, moving the AP initiation site away from the soma along the axon reduces the upper limit of the lateral current shaping the AP initiation dynamics. As for our case where *x*_Na_ is set to 40*μ*m, some upstroke dynamics was retained even when the soma was extremely large. The population encoding ability in the large soma model limit seems bounded by the sodium activation dynamics at the initiation site.

In Fig 11, we analyzed the electrotonic filtering and AP initiation dynamics of Brette’s models with different soma sizes. As a comparison, we also present the analysis for Brette’s models with high voltage sensitivity of the sodium activation function. We will show that the enhanced encoding ability seen in Fig 10 originates from the acceleration of axonal AP initiation dynamics.

**Figure 11.**
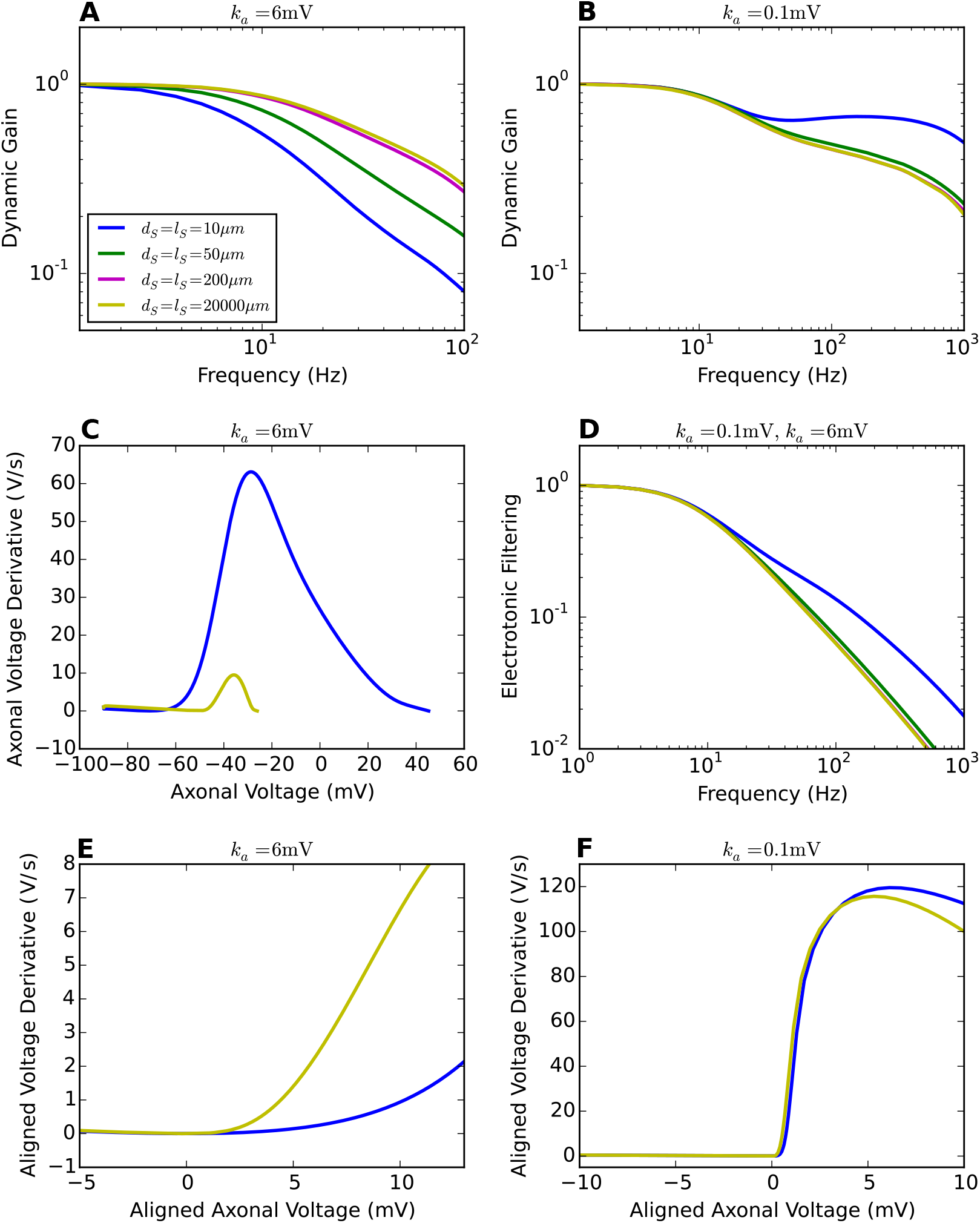
Electrotonic filtering and AP initiation dynamics for different soma sizes. **A** and **B** Dynamic gain functions of Brette’s model for a variety of soma sizes. *k_a_* is set to 6mV and 0.1mV respectively. *x_Na_* = 40*μ*m. *τ* = 5ms. Legend panel in **A**. **C** AP initiation dynamics seen at the axon with *k_a_* = 6mV. **D** Electrotonic filtering for different soma sizes. The passive neuron models are identical for both *k_a_* values. **E** and **F** Axonal AP initiation dynamics with local minima aligned at (0mV, 0mV/ms).

An increase in soma size changes the input encoding in two ways. The filter from input current to voltage at the initiation site is changing and an increased lateral current drains more of the depolarizing sodium current. The electrotonic filtering, shown in Fig 11**D**, decays faster with a bigger soma. This effect dominates for the steep sodium activation curve (*k_a_* = 0.1mV) and underlies the reduction in dynamic gain in Fig 11**B**. The second effect, the increased current sink of the larger soma, indeed led to a decreased AP initiation speed, but while this was almost negligible for *k_a_* = 0.1mV, it was very large for the conventional sodium activation curve (*k_a_* = 6mV)in Fig 11**C**. As a result, the voltage at which sodium currents overcome current drainage by leak and lateral current is shifted by more than 10mV to a region where the sodium activation curve is steeper. For better comparison, Fig **11E** aligns the local minima of two axonal AP initiation dynamics. The AP initiation dynamics are slower and smoother for the small soma. In this sense, the enormous soma size accelerates the initial AP initiation, creating a steeper axonal AP waveform and thereby an enhanced dynamic gain in the high frequency region.

## Discussion

Our study demonstrates that the experimentally observed high bandwidth encoding of cortical neurons is critically influenced by the properties of the ion channels in the AIS, and further modulated by the electrotonic structure of axo-somatic and somato-dendritic compartments. We used the dynamic gain function for an extensive comparison of model variants, designed to test the resistive coupling hypothesis ***Brette (2013***); ***Telenczuk et al. (2017***) and the impact of AIS ion channels. By choosing the same, well-defined working points for all models, we eliminated free parameters and assured unbiased comparison. Resistive coupling ***Goethals and Brette (2020***), also called compartmentalization ***Brette (2013***), is not sufficient to account for the ultrafast population coding. We found the dynamic gain function to be weakly influenced by the separation between AP initiation site and soma, as was the initial slope of the AP phase plot at the initiation site. A weak impact on AP initiation and population encoding was also observed when we changed the sodium conductance amplitude or the soma size. Even extreme choices for these parameters were insufficient to induce a high bandwidth of the dynamic gain function. An increased voltage sensitivity of sodium channel activation, however, substantially enhanced AP initiation speed and population encoding ability. Our results indicate that the key determinant of population encoding ability is the effective voltage dependence of AP initiating ion current, subsumed here into a single sodium channel type.

The close relation between the voltage dependence of AP initiation currents and the encoding bandwidth is consistent with previous theoretical studies. For model currents with exponential ***Fourcaud-Trocmé et al. (2003***) or piecewise-linear voltage dependence ***Wei and Wolf (2011***), increases in the voltage dependence substantially improved the encoding of high frequency input components. The low cut-off frequency that we report here for the model from ***Brette (2013***) with standard voltage sensitivity resemble the results of single compartment conductance-based models ***Huang et al. (2012***); ***Fourcaud-Trocmé et al. (2003***) in which a similar voltage dependence was used. Encoding of neurons with an infinite voltage sensitivity, represented by the fixed AP threshold in integrate and fire models, was first considered by ***Knight** (**1972a***) and more extensively studied by ***Brunel et al. (2001***). We closely reproduce its encoding behavior when we increase the axonal current’s voltage sensitivity. A remaining difference is the finite bandwidth, with is plausibly caused by the finite activation speed of our model, *τ_act_* = 0.1ms as compared to the instantaneous activation in a LIF model.

We reporta modulatory impact of neuron morphology, which is consistent with previous studies ***Eyal et al. (2014***); ***Ostojic et al. (2015***); ***Verbist et al. (2020***). The neurons electrotonic structure affects the effective impedance transforming fluctuating current input into voltage at the initiation site. ***Eyal et al. (2014***); ***Ostojic et al. (2015***); ***Verbist et al. (2020***) showed that for current injections proximal to the initiation site, the impedance of a large somato-dendritic compartment causes an apparent boosting of high-frequency components. On the other hand, an increasing distance between injection site and initiation site can attenuate high-frequency components, as we show in figure 8. For simple morphologies, such passive effects can be understood analytically ***Aspart et al. (2016***). Our analysis reveals another effect of the electrotonic structure, mediated by an altered AP initiation dynamics. An increased somatic current sink drastically reduces the depolarizing currents available at the initiation site. However, the voltage threshold for AP initiation is also strongly shifted towards the voltage at which the ion currents attain their peak voltage dependence (Fig 11). These two effects have opposing impact on the activation dynamics at threshold and consequentially, the combined effects of increased somato-dendritic size can reduce the bandwidth, as in the case of high voltage sensitivity *k_a_* = 0.1mV, or increased the bandwidth, as in the case of a low voltage sensitivity *k_a_* = 6mV, similar to the waveform AP waveform change reported by ***Eyal et al. (2014***). While several of our findings have been reported before in different models and with different emphasis, our study offers the first exhaustive comparison of models, preserving an invariant firing rate and ISI distribution under all parameter variations. It thus provides an unambiguous dissection of the different passive and active effects.

Over the past decade, the simple picture of the initial axon as a high channel density focus in an otherwise uniform axonal membrane has been superseded by findings of an ever more intricate nanoscale organisation of the AIS. Its structured by the interplay of cell-adhesion, cytoskeletal and scaffolding proteins and nanoscale ion channel clusters ***Xu et al. (2013***); ***Jones et al. (2014***); ***Ho et al. (2014***); ***Leterrier et al. (2015***). Dynamic gain studies consistently suggest that the AIS molecular nano-architecture is under substantial selective pressure to equip the population of hundreds to thousands of cells to which a particular neurons belongs, with a high encoding bandwidth ***Ilin et al. (2013)**; **Lazarov et al. (2018***); ***Revah et al. (2019***). From this perspective, the AIS appears as an intricately structured service organelle that is built and maintained by the individual neuron but fulfills its essential function only at the higher level of populations and neuronal circuits. The population coding speed hinges on the voltage sensitivity of the AP initiation current, and is modulated by the electrotonic structure of the neuron. In real neurons, more than one ion channel type contributes to the AP initiation dynamics and a pioneering study has already shown that not only Na currents, but also Kv1 potassium currents, activating around AP threshold, improved AP locking to high input frequencies ***Higgs and Spain (2011***). The critical importance of axonal conductances is further supported by observations of reduced encoding bandwidth following local manipulations of axonal conductances through pharmacological intervention or mutation of the axon cytoskeleton ***Ilin et al. (2013***); ***Lazarov et al. (2018***).

Systematic studies connecting AIS molecular architecture, to theory-driven, statistical analysis of population encoding performance are required to understand the impact of AIS molecular design, plasticity and pathology on population level information processing. Dynamic gain measurements are currently the most sensitive tool for that such an approach, as demonstrated by recent experimental and theoretical study ***Lazarov et al. (2018***); ***Verbist et al. (2020***). However, it is crucial that the comparison across neurons is performed systematically and without bias. Specifically, the AP trains evoked in such experiments should have the same overall statistics, because the dynamic gain function of a neuron depends on the operating point it is tested at. Here, we fixed the firing rate and the CV of the ISI distribution, following ***Vilela and Lindner (2009***). Such standardization assures that the firing patterns of different simulations have similar entropy, highlighting the actual population encoding differences across models.

In our dynamic gain analysis, we observed a clear impact of the voltage dependence of the axonal currents on the AP initiation dynamics and the bandwidth of population encoding. Several experimental studies have shown that the AIS can undergo plastic changes when their activity levels are changed ***Kuba et al. (2006, 2010, 2015***); ***Grubb and Burrone (2010***); ***Grubb et al. (2011***); ***Lezmy et al. (2017***); ***Wefelmeyer et al. (2015***). In the auditory system, deprivation of external input not only increased the length of AIS and sodium channel densities ***Kuba et al. (2010***) but also changed the dominant potassium channel type ***Kuba et al. (2015***), which enhanced the neuronal excitability. In response to increased stimulation, however, excitatory neurons in different systems were found to become less excitable and to shift their axonal ion channels away from the soma by several micrometers ***Grubb and Burrone (2010***); ***Lezmy et al. (2017***); ***Wefelmeyer et al. (2015***). In the simple model that we studied here, such smalls shifts did not lead to a pronounced change in dynamic gain or excitability, however, the study by Kuba et al. ***Kuba et al. (2015***) raises the intriguing possibility that AIS plasticity also affects the composition of the axonal ion channels and thereby the dynamic gain ***Lazarov et al. (2018***). Post-translational modifications of ion channels through enzymes or modulation through auxiliary subunits are expected to impact dynamic gain within minutes or hours. Studies of AIS plasticity should therefore be complemented by measurement of the dynamic gain. A particularly drastic change of operating conditions, transient hypoxia and spreading depolarization has recently been shown to lead to a disruption of the AIS organization, and in this case the bandwidth of the dynamic gain was indeed substantially reduced ***Revah et al. (2019***). While AIS plasticity can adjust neuronal excitability in a relatively short period of time, the dendrite morphology might impact the dynamic gain on developmental time scales or cause differences between species ***?Goriounova et al. (2018***); ***Mohan et al. (2015***) or individuals affected by AIS pathologies ***Wang et al. (2018***). Our dynamic gain analysis of Brette’s model provides an approach and baseline for analyzing different neuronal populations, mutants and manipulations as well as multi-compartment models. This analysis approach can be generalized to study how neuron morphology and axonal conductances contribute to the dynamic gain in more complicated scenarios.

## Methods and Materials

### Model

All simulations were performed with NEURON 7.3 ***Carnevale and Hines (2006***) as a module for Python 2.6. We used a ball-and-stick neuron model composed of a soma and an axon. All parameters were identical to the model, the equivalence of the models was confirmed by reproduction of published results (see Fig 1**A**). Here, the somatic compartment was modelled as a cylinder with equal length and diameter, which has the same effective capacitance as the sperical soma in ***Brette (2013***). The soma diameter was *d_s_* = 50*μ*m and its length was *l_s_* = 50*μ*m. The axon had diameter *d_A_* = 1*μ*m and length *l_A_* = 600*μ*m. Passive properties of the soma and axon are listed below. Axial resistance was *R_a_* = 150Ωcm; specific membrane capacitance was *c_m_* = 0.75*μ*F/cm^2^ and specific membrane resistance was *R_m_* = 30000Ωcm^2^ with leak reversal potential of *E_L_* = −75mV. The only voltage dependent conductance in the model represents sodium current at a single location, at a distance away from the soma. This was modeled as a NEURON point process. This sodium current has a Boltzmann activation curve similar to experimental observations ***Schmidt-Hieber and Bischofberger (2010)*** (*V*_1/2_ = −40mV *k_a_* = 6mV). This was combined with a voltage independent activation and deactivation time constant of *τ_m_* = 0.1ms. As in ***Brette (2013***), sodium channel inactivation was ignored. The sodium peak conductance was 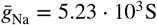. The reversal potential of the sodium current was *E_Na_* = 60mV. In a subset of simulations, the voltage dependence of the sodium current was changed as noted in the results.

The neuron model was thus defined by the following system of differential equations.

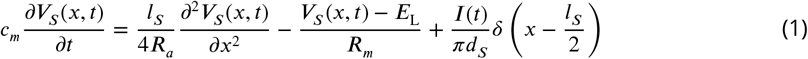

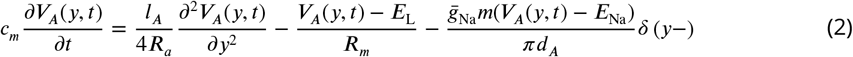

where

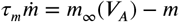

and

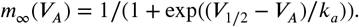

The voltage at the soma was denoted *V_s_*(*x*, *t*) with *x* ∈ [0, *l_s_*]. The voltage in the axon was denoted *V_A_*(*y*,*t*) with *y* ∈ [0, *l_A_*]. The stimulus current was injected in the middle of the soma, and denoted *I*(*t*). *δ*(·) was the Dirac delta function.

The boundary conditions of Eq (1) and (2) were given as 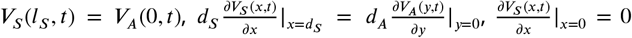, and 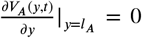. The first two conditions implied the continuity of voltage and current at the connecting point of soma and axon. The other two conditions represented the sealed ends of the neuron model.

### Simulations

To drive the neuron model, we injected Gaussian colored noise currents *I*(*t*) into the soma as in prior experimental and theoretical studies ***Köndgen et al. (2008***); ***Boucsein et al. (2009***); ***Tchumatchenko and Wolf (2011)**; **Higgs and Spain (2009, 2011***); ***Ilin et al. (2013***); **?**. We recorded the voltage at the position of the sodium channels in the axon and in the middle of soma. We refer to these as axonal and somatic voltage. We used the default backward Euler method in NEURON to integrate Eq (1) and (2) with a time step of Δt = 25*μ*s and a spatial grid of Δx = 1*μ*m.

An AP was detected, when the axonal voltage reached a detection threshold. For each variant of the model, this detection threshold was chosen as the axonal voltage at which the AP evolved most rapidly, i.e. the point of maximal rate of voltage rise. In this way, the AP detection time was least influenced by the ongoing random current fluctuations.

Because the neuron model does not feature sodium channel inactivation nor repolarizing voltage activated currents, we terminated each AP by a forced reset. Specifically we globally reset the membrane voltage to *E_L_* and the sodium gating variable *m* to *m*_∞_(*E*_L_). The reset is performed upon crossing the reset threshold, which is chosen such that on average reset occurs 2ms after the AP detection threshold was crossed.

We used an Ornstein-Uhlenbeck (OU) process ***Tuckwell (1989***); ***Gillespie (1996***) to create the current stimuli, which are characterized by their correlation time *τ*, standard deviation *σ* and mean current *μ*:

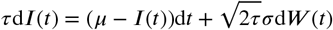

where *W*(*t*) denotes a Wiener process with zero mean and unit variance. A given set of mean and standard deviation results in AP sequences that can be characterized by the average firing rate *v* and the CV of the ISI distribution. Changes in the input parameter do change the operating point (*v*,*CV_ISI_*) of the neuron model, and thereby also its dynamic gain function. Therefore, we compare different model variants at the same operating point: *v* = 5±0.25Hz and *CV_ISI_* = 0.85±0.05, representing in the fluctuation-driven regime in which cortical pyramidal neurons operate *in vivo*. For a subset of simulations, different operating points were chosen, as stated in the results. The OU parameters *σ* and *μ*t hat drive the model to this operating point were determined in independent simulation runs.

For studies of the impact of sodium peak conductance on population encoding, the voltage values were reset to −90mV after an AP to maintain type 1 excitability throughout the whole range of sodium peak conductances.

To compare the AP waveforms for different model variants, we injected each model with the constant input that generated the target firing rate. The local minima of phase plots were aligned at (0mV, 0mV/ms).

### Linear response, dynamic gain and electrotonic filtering

To compute the dynamic gain of a neuron model, that is, its linear response function from the injected current to the firing rate, we followed ***Higgs and Spain (2009***) as outlined below. The linear response function was given by

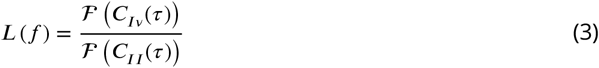

where 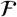 denotes the Fourier transformation, *C_Iv_* and *C_II_* the input-output correlation and input auto-correlation functions, respectively. The Fourier transform of the input auto-correlation equals the power spectral density of the input, according to the Wiener-Khinchin theorem. For an Ornstein-Uhlenbeck process, the power spectral density is 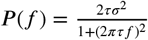.

To compute the input output correlation function we made use of the following equality

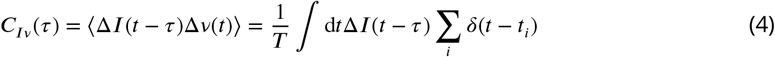

where *δ*(*t* - *t_i_*) is the Dirac delta function, Δ*v* and Δ*I* denote deviations from mean firing rate and mean input respectively. *T* is the simulation time. Thus *C_Iv_*(*τ*) is the spike-triggered average current (*STAļ*) multiplied with the firing rate (*v*). The time window to compute the (*STA_I_*) was chosen such that correlations decayed within the time window, which was fulfilled by a window of size 800ms centered about the spike time. We then computed the Fourier transform and applied a bank of Gaussian filters to the complex-valued Fourier transform components of the STAto de-noise as proposed in ***Higgs and Spain (2009***).

The dynamic gain was calculated from 20000 trials of numerical simulations of the stochastic current driven model described above. Each trial represents 20.5s of real time (with 0.5s burn-in time and 20s of actual recording). From the 20000 pairs of input current samples and output AP trains we determined the *STA_I_* through Eq (4), calculated the linear response function with Eq (3) and finally obtained the dynamic gain as its absolute value *G*(*f*) = |*L*(*f*)|.

For each model variant we determined a significance threshold curve according to the zero-hypothesis that the AP times are independent from the input waveform and hence the dynamic gain is zero. The zero hypothesis was realized by cyclically shifting all AP times of a given trial by the same, randomly chosen interval between 1 and 19 seconds. From all 20000 pairs of input current and shifted AP times, a dynamic gain curve was calculated. This was repeated 500 times for different random time shifts. The 95 percentile of these curves determined the significance threshold curve.Only significant parts of the dynamic gain functions are shown in the following figures. For the bootstrap confidence interval, we used the 20000 individual simulations to calculate 400 *STA_I_* curves from 50 simulations each. We then bootstrap re-sampled the grand average *STA_I_* from these 400 STA estimates for 1000 times and obtained the corresponding dynamic gain curves. The 95 percent confidence intervals from these bootstraps are shown for each dynamic gain curve, typically they are smaller than the line width.

We assessed the electrotonic filtering of stimuli transmitted from soma to the AP initiation site in purely passive neuron models, models, sodium peak conductance 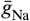 was set to 0. We injected the soma with sinusoidal stimuli of 1Hz to 1000Hz, and recorded voltage fluctuations along the axon. The electrotonic filtering function was calculated as the amplitude ratio of the Fourier transform of output voltage fluctuations to the Fourier transform of the input stimuli at corresponding frequencies.

## Acknowledgments

We thank Barbara Feulner and Rainer Engelken for fruitful discussions. This work was supported by the China Scholarship Council (to C.Z.), the United States National Institute of Health (NIH) BRAIN Initiative Theory Grant 1R01-EB022872 (to D.H.), and NIH grant 5R01-NS099375 (to D.H.), the German Federal Ministry for Education and Research (BMBF) under grant no. 01GQ1005B (to A.P., T.G., F.W. and D.B.), through CRC 889 and NeuroNex Working Memory by the Deutsche Forschungsgemeinschaft and by the VolkswagenStiftung under grant no. ZN2632 (to F.W.).

## Notes

### Competing Interest Statement

The authors have declared no competing interest.

